# Comparing alternative pipelines for cross-platform microarray gene expression data integration with RNA-seq data in breast cancer

**DOI:** 10.1101/059600

**Authors:** Alina Frolova, Vladyslav Bondarenko, Maria Obolenska

## Abstract

**Background:** According to major public repositories statistics an overwhelming majority of the existing and newly uploaded data originates from microarray experiments. Unfortunately, the potential of this data to bring new insights is limited by the effects of individual study-specific biases due to small number of biological samples. Increasing sample size by direct microarray data integration increases the statistical power to obtain a more precise estimate of gene expression in a population of individuals resulting in lower false discovery rates. However, despite numerous recommendations for gene expression data integration, there is a lack of a systematic comparison of different processing approaches aimed to asses microarray platforms diversity and ambiguous probesets to genes correspondence, leading to low number of studies applying integration.

**Results:** Here, we investigated five different approaches of the microarrays data processing in comparison with RNA-seq data on breast cancer samples. We aimed to evaluate different probesets annotations as well as different procedures of choosing between probesets mapped to the same gene. We show that pipelines rankings are mostly preserved across Affymetrix and Illumina platforms. BrainArray approach based on updated annotation and redesigned probesets definition and choosing probeset with the maximum average signal across the samples have best correlation with RNA-seq, while averaging probesets signals as well as scoring the quality of probes sequences mapping to the transcripts of the targeted gene have worse correlation. Finally, randomly selecting probeset among probesets mapped to the same gene significantly decreases the correlation with RNA-seq.

**Conclusion:** We show that methods, which rely on actual probesets signal intensities, are advantageous to methods considering biological characteristics of the probes sequences only and that cross-platform integration of datasets improves correlation with the RNA-seq data. We consider the results obtained in this paper contributive to the integrative analysis as a worthwhile alternative to the classical meta-analysis of the multiple gene expression datasets.

## Background

Rapid advances in high-throughput technologies have led to the generation of a large amount of genome-wide data that is widely used in biomedicine, and resulted in a considerable progress in the human disease biomarkers discovery and therapy [1]. Major experimental data repositories such as ArrayExpress and Gene Expression Omnibus (GEO) report exponential growth of the fraction of next-generation sequencing data among their new submissions over the last few years. Nevertheless, the total number of assays associated with such experiments is still very low in comparison with microarray-based experiments [2, 3]. The latter is mostly a result of microarrays being a more accessible tool for transcriptomic analysis of a large number of clinical samples with lower input requirements in contrast to next-generation sequencing, which is more expensive along with its more time and resource consuming analysis and data storage [4, 5]. Also, the microarray gene expression data often remains to be the only one source of knowledge about expressed transcripts in particular cell types and tissues in health and disease. But in order to make sound conclusions about the tested biological hypotheses, the researchers require sufficient number of samples, which are not always available from the individual studies. Increasing sample size by direct data integration increases the statistical power to obtain a more precise estimate of gene expression in a population of individuals and to reduce the effects of individual study-specific biases [1]. However, such integration implies that the data is typically originated from different studies and microarray platforms, leading to a number of technical issues.

One of such issues is an accurate mapping of the microarray probes to their targeted sequences as the information about the probes annotation is sometimes incomplete and outdated. For example, it was reported that over 25% of Illumina chip probes could be also mapped to genes other than those in the manufacturers annotation, which emphasizes importance of the most up-to-date annotation usage [6]. To this end, the typical option is to use Bioconductor libraries, that provide annotations for a variety of microarray platforms and are updated twice a year [7, 8]. The more advanced way of mapping probe sequences to the targeted genes or transcripts is using sequence alignment tools [9, 10]. For instance, researchers may use tools that are specifically designed for the probes re-annotation, such as Re-MOAT or Re-Annotator [11, 6] or utilize BLAST [12] to map probes from different platforms to the up-to-date transcriptome sequences [8, 13]

Another issue is the ambiguous correspondence between probesets and their targeted genes. Here, we focus on two major microarray gene expression chip platforms, Affymetrix and Illumina, since both of them are most widely used and show consistent results in spite of different probe design and structure [14]. Affymetrix chips carry short 25-mer oligonucleotides, called probes, where a collection of target-specific probes with different sequences is called a probeset. Thus, the summarized signal intensity value of probes in a probeset corresponds to expression level of a particular gene. On the contrary, Illumina chips are based on silica beads covered with long 50-mer oligonucleotides, where each bead is represented by, on average, 30 instances within the array, providing an internal technical replication [15]. For simplicity and coherence, we are going to use term ‘probeset’ when referring to signal intensity values summarized from the Illumina beads of the same instance. However, no array design provides an absolute absence of multiple probesets mapped to the same genes [13]. Moreover, some probes in the probesets of Affymetrix arrays can be mapped to different genes or even non-coding regions [16, 1]. Since integrating microarray data from different platforms using gene identifiers requires one-to-one (unique) correspondence between probesets and genes, there have been proposed several solutions in this field.

For example, the problem of choosing the best probeset within a group of probesets mapped to the same gene can be solved by assigning scores to each probeset that reflect some of their important characteristics such as specificity or sensitivity to a given DNA sequence [16, 17]. Other studies suggested averaging signal intensities of probesets mapped to the same gene [18, 13], selecting probeset with the highest interquartile range of the signal intensity [19], taking median intensity value of the probesets [20, 21], choosing probeset with the highest average intensity [22, 23, 24] or choosing a random probeset [25]. Finally, there is also a solution based on the redesign of the manufacturers probesets in a way to assemble all probes that are specific to the same gene into a single probeset [9, 10, 26, 27, 28]. Suchwise, this procedure produces a list of probesets, which have one-to-one correspondence with a list of genes. However, despite the variety of proposed solutions, there are no studies where these strategies have been systematically evaluated and compared on data from different microarray platforms.

The other issue related to microarray data integration is a cross-platform normalization, which deals with the batch effect. The latter is a collection of systematic errors, which appear when samples are processed in multiple batches, or, in other words, it is a cumulative error introduced by time and place-dependent experimental variations [29]. Although a batch effect can be reduced by careful experimental design, it cannot be eliminated unless the whole study is done in a single batch. Among different cross-platform normalization methods, ComBat (or Empirical Bayes method), is recommended when merging datasets as it generates biologically meaningful results with improved statistical power [13, 30, 31, 24].

Despite numerous recommendations for gene expression data integration, the most recent comprehensive systematic literature overview of the studies applying microarray integration methods, revealed that only 27% of the studies utilized direct microarray data integration, and this subset of studies was mostly performed on the same platforms [1, 32]. We suggest that this is caused by the lack of a systematic evaluation of the impact of different pipelines of the probesets annotation and selection. Therefore, here we validate different microarray processing approaches by comparing them with the RNA-seq data as it was shown to be correlated well with microarray data from different platforms [23, 5, 17]. Additionally, we consider RNA-seq being a good reference for validation, because it outperforms microarrays by several factors: provides a wider dynamic range for measuring transcripts abundance, and has lower background and higher resolution [25, 33]. As to our test system choice, breast carcinomas have been extensively profiled by high-throughput technologies for over a decade, and a large amount of data from prognostic and/or treatment predictive gene expression studies is available in public repositories and was already used in a number of various studies about the microarrays analysis [27, 13] as well as in the studies comparing microarrays and RNA-Seq [17].

Thus, in our study we focus on the problems related to the probesets reannotation, getting a unique probeset-to-gene correspondence, and aim to explore how different approaches to microarray cross-platform data integration affect results of the gene expression studies of human breast cancer through the comparison with RNA-seq data. Namely, we show that methods, which rely on actual probesets signal intensities, are advantageous over the methods considering the biological characteristics of the probes sequences only and that combining datasets improves correlation with the RNA-seq data.

## Materials and Methods

### Datasets description

We considered three very popular microarray platforms: Affymetrix HG-U133 Plus 2.0, Affymetrix HuGene 1.0 ST, and Illumina HumanHT-12 v4.0. Three raw Affymetrix microarray gene expression datasets, one non-normalized summary-level Illumina chip dataset, and one raw RNA-seq dataset were retrieved from the GEO repository (Table 1). All samples represent tumor tissues collected from the patients with a preoperative diagnosis of primary invasive breast cancer. Additionally, we obtained raw bead-level Illumina HumanHT-12 v4.0 dataset from the authors, which was used only for Illumina arrays normalization methods testing. This dataset contains samples from breast cancer associated fibroblasts and is available from the GEO in a summary-level format under the GSE37614 identifier [34].

In order to assign cancer subtype to each sample we used scheme from Onitilo et al study, where breast cancer is classified into four groups based on immunohistochemistry (IHC) positive (+) and/or negative (−) profile of estrogen/progesterone receptors (ER/PR) and human epidermal growth factor receptor 2 (Her2) expression. The IHC classification correlates with intrinsic (molecular) gene expression microarray categorization: **ER/PR**+, **Her2**+ with Luminal B; **ER/PR**+, **Her2-** with Luminal A; **ER/PR-**, **Her2**+ with HER2 and **ER/PR-**, **Her2-** – with basal-like tumors [35].

**ER/PR-**, **Her2-** IHC subtype, also known as triple negative breast cancer (**TNBC**), shows 80% concordance with its intrinsic “equivalent” (basal-like breast cancer) and is distinctively clustered apart other molecular subtypes regardless of the cancer subtyping scheme and classification methods [36, 37]. As pooling datasets with a high degree of heterogeneity might complicate subsequent analysis, we used only TNBC samples, which are mostly of histological grade III (Table 2), when comparing microarrays and RNA-seq data. The whole microarray datasets were used for quality assessment.

**Table 1.** Description of the datasets used in the study. Cancer molecular subtypes were determined based on the immunohistochemistry classification and its correlation with intrinsic schema.

| Study GEO ID<br>[reference] | Platform | TNBC | lumA | lumB | Her2 |
| --- | --- | --- | --- | --- | --- |
| GSE65194 [38] | Affymetrix HG-U133 Plus 2.0 | 55 | 29 | 30 | 39 |
| GSE19615 [39] | Affymetrix HG-U133 Plus 2.0 | 31 | 25 | 30 | 13 |
| GSE58644 [40] | Affymetrix HuGene 1.0 ST | 50 | 206 | 38 | 20 |
| GSE60785 [37] | Illumina HumanHT-12 v4.0 | 13 | 33 | 3 | 1 |
| GSE58135 [41] | Illumina HiSeq 2000 | 42 | – | – | – |

### Study design

The schematic representation of workflow is given in Fig. 1. Hereinafter, we use “one-to-one” term when refer to genes or probesets, which have unique correspondence, “one-to-many genes” term – to genes with multiple probesets, and “many-to-one probesets” term – to the group of the probesets mapped to the same gene.

**Figure 1.**
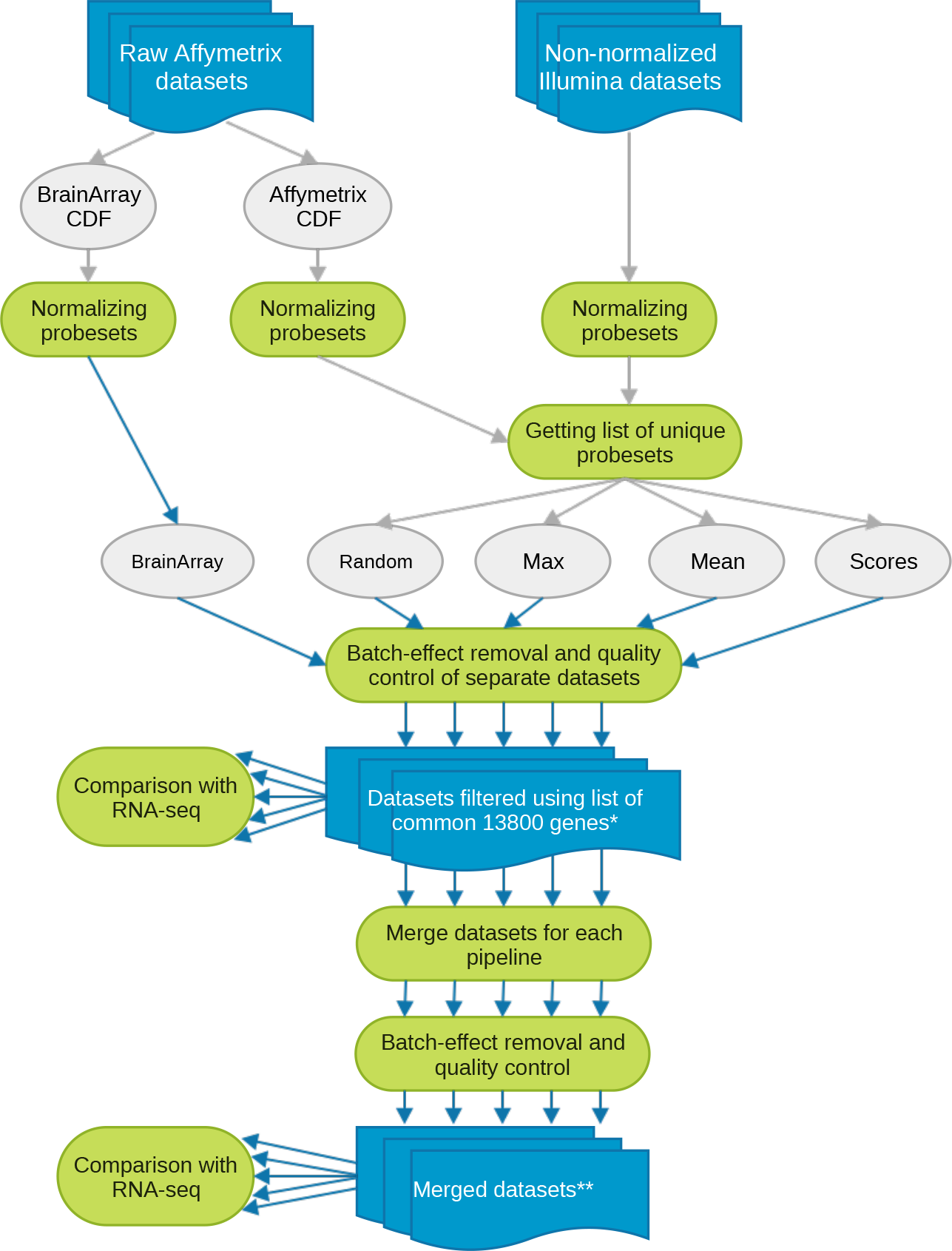
Data processing workflow. Each box corresponds to a separate step in the data analysis pipeline; blue lines denote different pipelines. * The number of datasets at this step is 19 – three datasets per Brainarray pipeline (not applicable for Illumina chip) and four datasets per other four pipelines. ** The number of datasets at this step is five – one dataset per each pipeline, since all the datasets within each pipeline were merged.

**Table 2.** The relation of tumors histological grade and cancer subtypes of breast cancer samples pulled from four microarray datasets.

| Breast cancer subtype | Tumor histological grade |  |  |
| --- | --- | --- | --- |
|  | I | II | III |
| TNBC | 1 | 8 | 137 |
| Luminal A | 65 | 132 | 95 |
| Luminal B | 9 | 24 | 68 |
| Her2 | 0 | 6 | 67 |

#### Microarrays data processing pipelines

**“Brainarray pipeline”** implies using Brainarray Chip Definition Files (CDF) version 19, where probes are reorganized into new probesets based on the latest genome and transcriptome information in a way to provide a unique correspondence between probesets and genes [42].

On the contrary, when analyzing Affymetrix datasets with native Affymetrix CDF and Illumina datasets we perform a key step to get unique genes-to-probesets list, which is required for the data merging. Namely, we chose three strategies: select random probeset among the many-to-one probesets – **“Random pipeline”** ; select the probeset with the maximum average signal intensity value across the samples – **“Max pipeline”** ; or take average signal intensity value of the many-to-one probesets – **“Mean pipeline”**. Also we implemented and tested two types of the Max pipeline: calculating maximum average intensity value across TNBC samples only, and calculating it across all samples in the dataset, i.e. including other cancer molecular subtypes (”Maxoverall” pipeline). Using Bioconductor annotations, we excluded probesets that simultaneously map to different genes.

In the **“Scores pipeline”**, many-to-one probesets are ranked according to “specificity” and “coverage” scores, where specificity designates mapping quality of probes sequences to the transcripts of the targeted gene, and coverage – potential to recognize all transcripts of the same gene, according to Li et al [16]. Namely, the probe was considered specific if it had a BLAST alignment score >32 for Affymetrix and >63.5 for Illumina chips; probesets which have less than a half of specific probes were excluded. Next, we defined the targeted gene of the probeset as a gene, which transcripts are specifically recognized with the majority of probes in the probeset. Finally, the **specificity score** of the probeset is defined as its probes average specificity. Whereas **coverage score** is defined as the number of transcripts of the targeted gene, which are specifically detected by the probes in the probeset, divided by the number of all transcripts of the targeted gene in the RefSeq annotation (Suppl. Fig. 1).

Thus, we processed four selected microarray datasets with five different pipelines, which resulted in 19 datasets as Brainarray pipeline is not applicable for Illumina chips. After selecting common genes based on Entrez IDs, we compared each dataset with RNA-seq data. Finally, we merged datasets within each pipeline and compared resulted datasets with RNA-seq data.

#### General data processing steps

Beadarray R package [43] was used to transform and normalize raw bead-level Illumina HumanHT-12 v4.0 dataset (GSE37614).

Following the scheme in Fig. 1, normalization of Affymetrix arrays was performed with the RMA method from affy package [44], while Illumina arrays were quantile normalized using lumi package [45]. Batch-effect removal was performed using ComBat method from the sva package [46] and quality control – using ArrayQualityMetrics package (Suppl. Fig. 2-7) [47].

Paired-end RNA-seq reads were mapped to the hg19 human genome with tophat2 [48] using RefSeq 67 human transcriptome annotation, following transcripts abundance calculation and normalization in FPKM (Fragments Per Kilobase of transcript per Million mapped reads) with cufflinks pipeline [49]. For further analysis, transcripts with FPKM over the threshold 0.01 were considered. Multiple transcripts per gene were eliminated by summing the expression values or by choosing maximum, although the resulting lists didn’t yield any substantial difference.

When comparing microarray and RNA-seq datasets mean expression values over the TNBC samples were considered.

All the programming was done using Bioconductor version 3.1 (BiocInstaller 1.18.5) and R version 3.2.3. The source code is available from github repository.

## Results

### Variation of the probesets intensity values across many-to-one probesets in different microarrays platforms

In each microarray platform one-to-many genes are mostly represented by 2–5 probesets, while the maximum, 37 probesets per gene, was identified in Illumina platform (Suppl. Fig. 8). In order to assess the signals variation of many-to-one probesets, we calculated standard deviation of probesets signal intensity values within each many-to-one group with respect to corresponding gene expression values from the RNA-seq data. Fig. 2 shows that the standard deviations grow as expression values of corresponding genes increase and reach its maximum level of around 6 in log2 scale independent of the platforms. Importantly, the observed growth is not the effect of the samples diversity within each dataset as we also compared the standard deviation of many-to-one probesets between the samples (instead of averaging probesets values across the samples), which showed no direct correlation with the increase of RNA-seq gene expression values (Suppl. Fig. 9). Therefore, choosing between many-to-one probesets can influence the results of the microarray data evaluation by introducing additional variation, especially for highly expressed genes.

We also compared two different datasets of the same Affymetrix HG-U133 Plus 2.0 platform without applying any additional probeset selection procedure as we could match the initial lists of probesets. Despite the fact that samples from different patients were used in those studies, a very high correlation was observed between the datasets. Meanwhile, probesets signal intensity values of Affymetrix HG-U133 Plus 2.0 and Affymetrix HuGene 1.0 ST platforms correlated worse, confirming that the major inconsistency between the studies is rather introduced by the usage of different microarray platforms than the variance between samples (Fig. 3) [14, 28].

**Figure 2.**
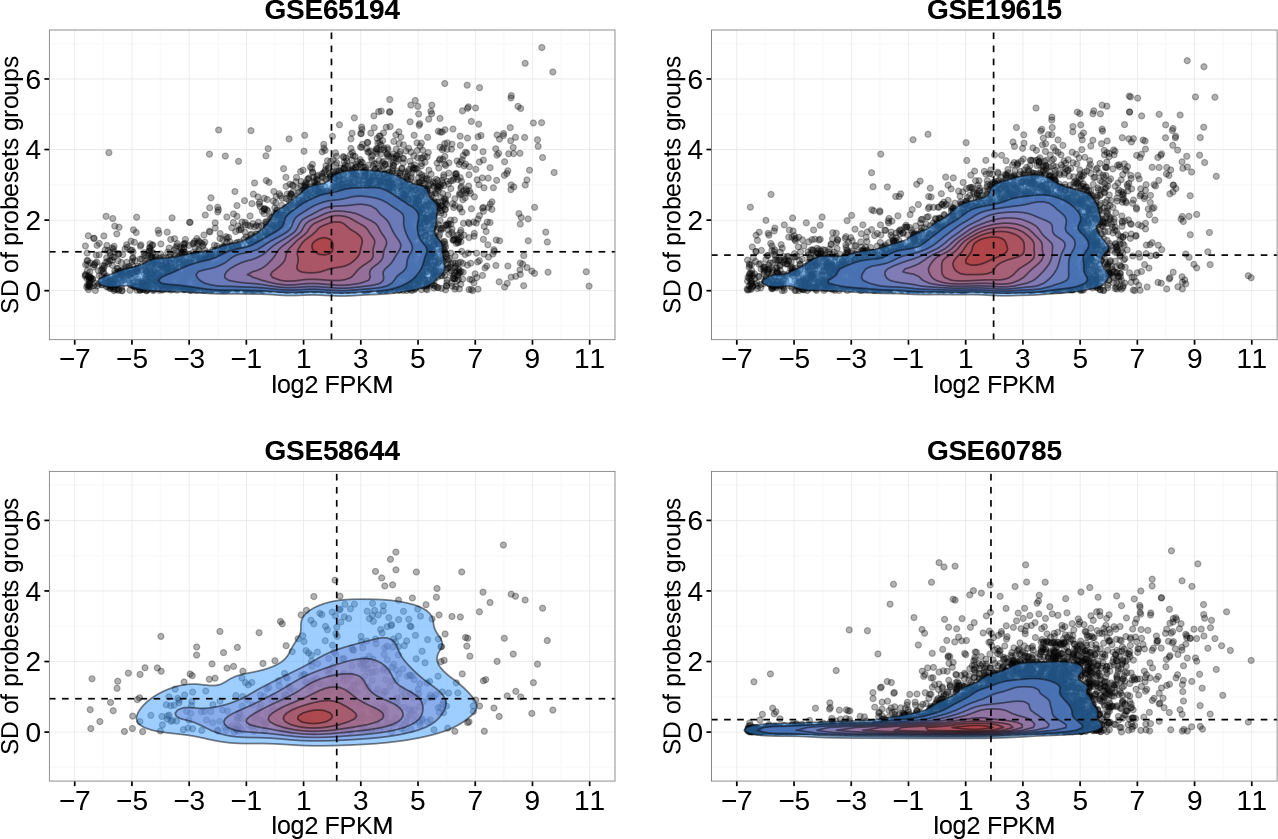
Variation of the probesets intensity values across many-to-one probesets in four microarray datasets versus corresponding genes values of RNA-seq. Each point on the graph refers to the standard deviation of the probesets intensity values across the group of probesets mapped to the same gene. The overall expression values range for GSE60785 dataset (Illumina chip) is 6-14 in log2 scale, while for the rest of datasets (Affymetrix arrays) – 2-14. Dashed black lines denote medians of the values distributions.

**Figure 3.**
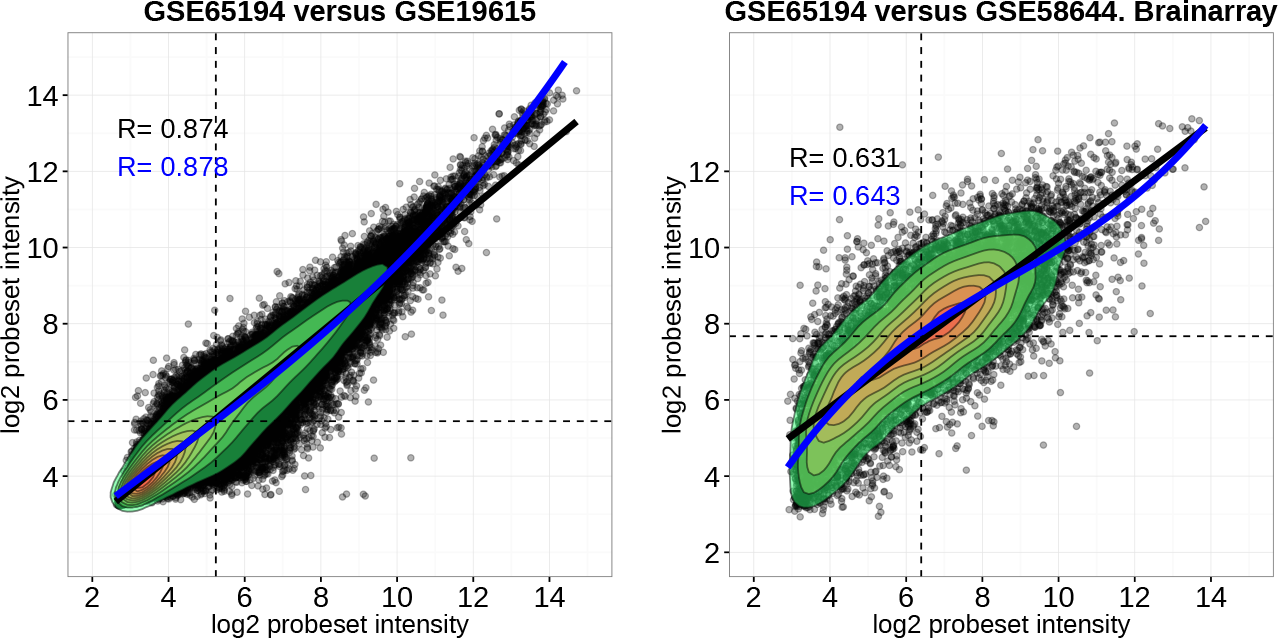
Expressional consistency between microarray data of different platforms. Left scatter plot depicts two datasets of the Affymetrix HG-U133 Plus 2.0 microarray platform on the entire range of the probesets; the right scatter plot illustrates Affymetrix HG-U133 Plus 2.0 platform in comparison with Affymetrix HuGene 1.0 ST using BrainArray annotation. Black dashed lines denote medians of the distributions; black and blue solid lines are fitted linear and cubic regression, respectively; R is a coefficient of determination.

### Microarray platforms characteristics according to different annotations

In order to compare Affymetrix HG-U133 Plus 2.0 and HuGene 1.0 ST, and Illumina HumanHT-12 v4.0 microarray platforms we calculated the ratio of uniquely represented genes versus genes represented by several probesets using three different probesets annotations: Bioconductor, Brainarray and Scores pipeline annotations (Table 3). Affymetrix HuGene 1.0 ST is the most specific microarray platform, having 2% and 13% of one-to-many genes according to Bioconductor and Scores annotations, respectively. Obviously, Brainarray annotation has 100% of unique genes as it was designed with this purpose. Score pipeline annotation is characterized by the smallest amount of annotated genes, because many of probesets didn’t pass the scoring procedure described in the previous section. Importantly, Table 3 shows that the number of genes left after intersecting genes lists between platforms within each annotation is notably decreased by nearly three thousand genes, which should be considered by researches when merging datasets from different platforms.

In the end, after intersecting the list of common 14246 genes for all the platforms and annotations with 23187 genes from RNA-seq dataset based on the gene EntrezIDs we have got 13800 common genes with 26% of one-to-one genes, which we used in further analysis.

**Table 3.**
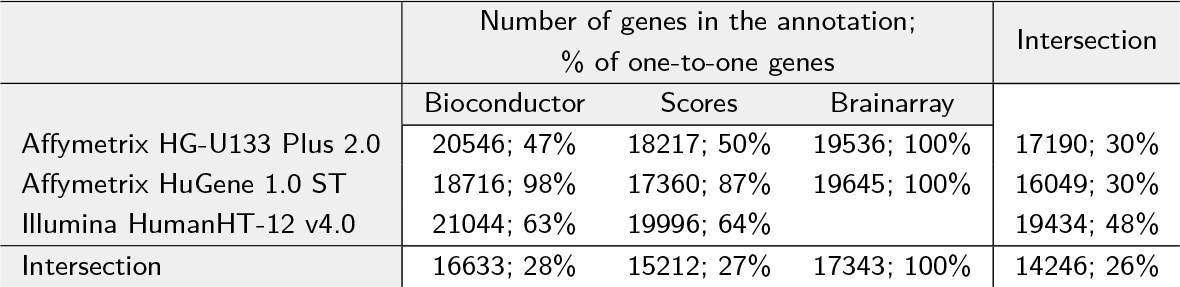
Total number of genes and the ratio of one-to-one among them in microarray platforms according to different annotations. Genes EntrezIDs are used for intersection between annotations/platforms genes lists. Each intersection decreases the number of one-to-one genes as they are supposed to be one-to-one for all the initial genes lists.

|  | Number of genes in the annotation;<br>% of one-to-one genes |  |  | Intersection |
| --- | --- | --- | --- | --- |
|  | Bioconductor | Scores | Brainarray |  |
| Affymetrix HG-U133 Plus 2.0 | 20546; 47% | 18217; 50% | 19536; 100% | 17190; 30% |
| Affymetrix HuGene 1.0 ST | 18716; 98% | 17360; 87% | 19645; 100% | 16049; 30% |
| Illumina HumanHT-12 v4.0 | 21044; 63% | 19996; 64% |  | 19434; 48% |
| Intersection | 16633; 28% | 15212; 27% | 17343; 100% | 14246; 26% |

### Comparison of microarray gene expression levels obtained along different pipelines with RNA-seq

Next, we compared gene expression levels between microarray datasets processed with different pipelines and RNA-seq data. Particularly, we looked at the three categories of genes: all 13800 common genes (Suppl. Fig. 10), genes that are represented by a unique probeset, or one-to-one genes (Suppl. Fig. 11), and genes that are represented by multiple probesets, or one-to-many genes (Suppl. Fig. 12). For each microarray platform we used one-to-one/one-to-many genes specific only for that platform (among 13800 genes).

Gene expression scatter plots shows non-linear trend between microarray data and RNA-seq especially observed in the range of low-expressed genes, which is a result of the extended dynamic range of the RNA-seq (or equivalently, the compressed dynamic range of microarrays) [33]. The latter is in agreement with RNA-seq considered being more advantageous over microarrays for detecting low-expressed genes [25, 23]. The type of a non-linear dependency between microarrays and RNA-seq as well as fitted curves share common traits with other studies [5].

We also considered coefficients of determination of linear and polynomial (cubic) regressions. Since microarray and RNA-seq data are not strictly linearly dependent, we wanted to see for which platforms and pipelines fitting polynomial regression gives better approximation, indicating that the data is less correlated linearly. On this basis, Affymetrix HuGene 1.0 ST platform differs even though it doesn’t have the highest R coefficients. First, it has wider dynamic range for low-expressed genes, second, the residuals and scale-location diagnostics plots shows that fitting cubic regression results in model overfitting in comparison to other platforms (Suppl. Fig. 13 and 14), which might indicate that HuGene 1.0 ST platform is more linearly correlated with RNA-seq than other platforms.

Another finding is that Illumina HumanHT-12 v4.0 platform shows the worst correlation with RNA-seq irrespective to processing methods, especially for the low-expressed genes that have expression in range 6-8 in a log2 scale (Fig. 4). The most relevant information we found was a comparison of Illumina HumanWG-6 v1 microarray data and AB SOLiD 4 System RNA-seq data. Particularly, Brau et al. compared the correlation of protein-coding transcripts from RNA-seq data with previously published microarray datasets of the platelet protein-coding transcriptome, showing rather low Spearman correlation [50]. The majority of Illumina platform datasets uploaded to GEO or ArrayExpress databases are summary-level data preprocessed with GenomeStudio software. However, the GenomeStudio option of subtracting the average of intensity signals in negative controls has been shown to be flawed on several occasions [51]. To ensure that we do not spot the flaws of data preprocessing, we used dataset with bead-level data (idat files) [34] to test different data transformation and background correction methods. In particular, we compared quantile normalization with log2 transfomation, vsn (variance stabilization and normalization [52]) and neqc (normexp background correction using control probes [53]) methods. The results did not yield any substantial difference for low-expressed genes, though the shape of the signal distributions slightly differed between the processing methods with neqc giving the highest R coefficients (Suppl. Fig. 15).

**Figure 4.**
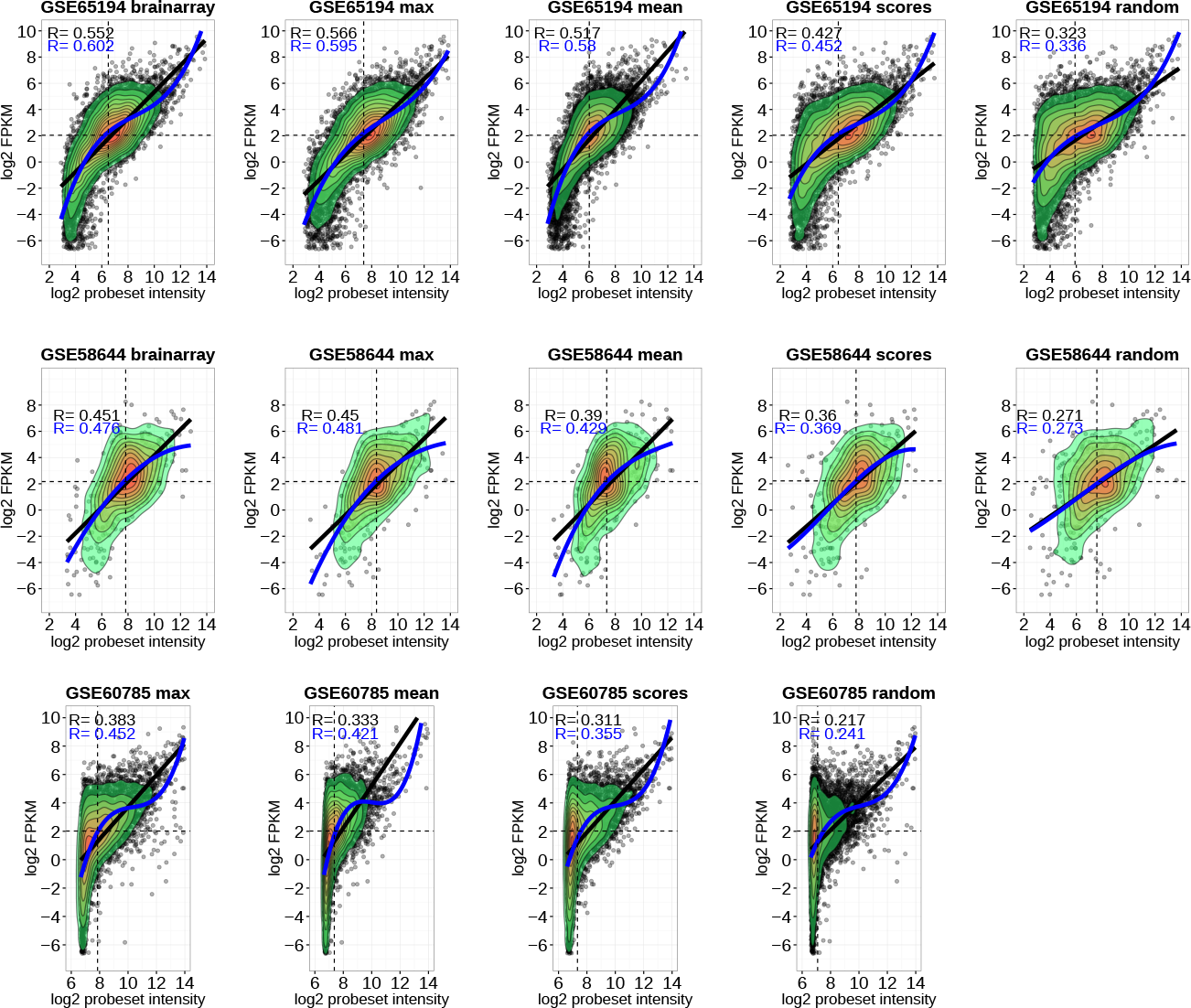
Comparison of the gene expression levels between RNA-Seq and microarray datasets on one-to-many genes. Log2 transformed FPKM values from RNA-Seq and log2 transformed microarray probesets intensity values were used for the scatterplots. Black dashed lines denote medians of the distributions, black and blue solid lines denote fitted linear and cubic regression, respectively; R is a coefficient of determination.

For the one-to-one category of genes (Suppl. Fig. 11), we did not observe any major difference between Bioconductor, BrainArray and Scores pipeline annotations for Affymetrix microarrays datasets, whereas for Illumina platform Scores showed better results than Bioconductor in terms of linear regression coefficient of determination.

Evidently, different methods of probeset selection from many-to-one probesets affect the correlation between microarrays and RNA-Seq data (Fig. 4). Particularly, ranking of the pipelines with respect to best concordance with RNA-Seq was reproducible on Affymetrix HG-U133 Plus 2.0 and Illumina datasets: 
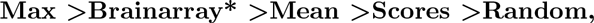

(where Brainarray pipeline is not available for Illumina datasets), whereas for Affymetrix HuGene 1.0 ST platform the ranking of the pipelines was the following:

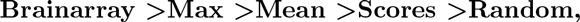

indicating that the best strategy of the probesets selection depends on the microarray platform. Notably, selecting random probesets gives the worst results in all the cases for all the platforms.

Comparing gene expression levels of one-to-many genes from microarrays their expression measured by RNA-seq using both Spearman and Pearson correlation resulted in a similar ranking of pipelines as using linear regression (Fig. 5). Namely, there is no substantial difference between Max and Brainarray pipelines correlation with RNA-seq, while Mean pipeline correlated better than Scores, and Scores better than Random. Maxoverall pipeline showed no substantial difference with usual Max pipeline, which is relevant in cases where one cannot define distinct group of samples before processing the data and therefore has to choose the probeset based on maximum average expression value across all the samples.

**Figure 5.**
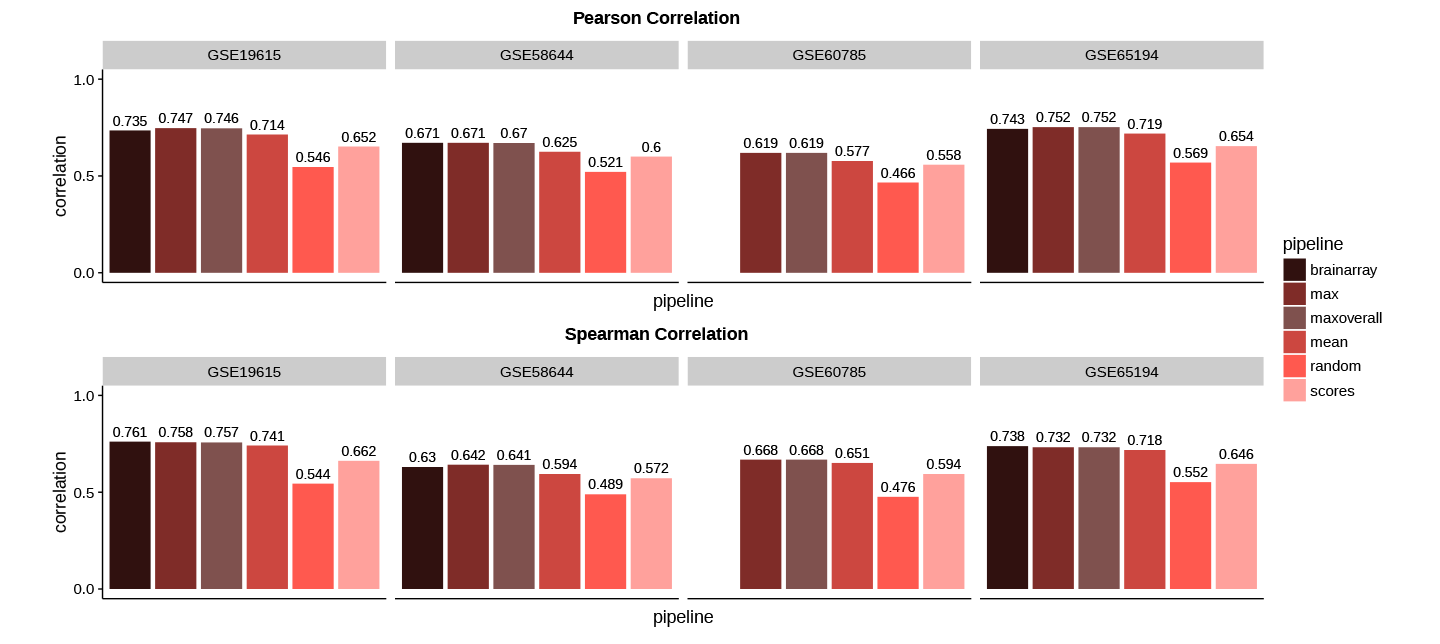
Pearson and Spearman correlation of the one-to-many genes expression levels between microarrays and RNA-seq data.

Correlations between expression values of one-to-one genes for Brainarray, Bioconductor and Scores pipelines probesets annotations do not differ substantially (Suppl. Fig. 16) being around 0.8. Additionally, when using the entire set of common 13800 genes it is evident that Illumina array has lower correlation with RNA-seq data than other platforms irrespective to the pipelines (Suppl. Fig. 17).

### Comparing merged microarray datasets with RNA-seq data

Finally, we combined microarray datasets within each pipeline and compared them with RNA-seq data using three groups of genes again: all, one-to-one and one-to-many. We found that combining datasets increased the correlation with RNA-seq for all the pipelines (Suppl. Fig. 18-20). Max and Brainarray pipelines perform almost similarly, while Mean >Scores >Random ordering still retains (Fig. 6).

**Figure 6.**
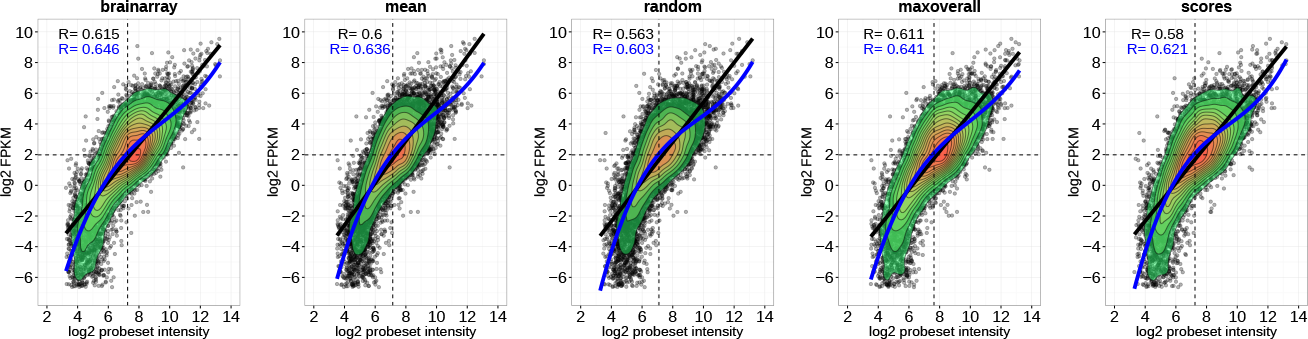
Comparison of the gene expression levels between RNA-seq and combined microarray datasets on one-to-many genes. Log2 transformed FPKM values from the RNA-Seq and log2 transformed microarray probesets intensity values were used for the scatterplots. Black dashed lines denote medians of distributions, black and blue solid lines denote fitted linear and cubic regression, respectively; R is a coefficient of determination..

### PCA plot of combined microarray dataset

We performed principal component analysis of the combined microarray dataset with respect to cancer subtypes. Fig. 7 confirms that there is a distinctive relation with the samples histological grade and subtypes clustering reported in previous studies [54], which is preserved after datasets processing and integration.

**Figure 7.**
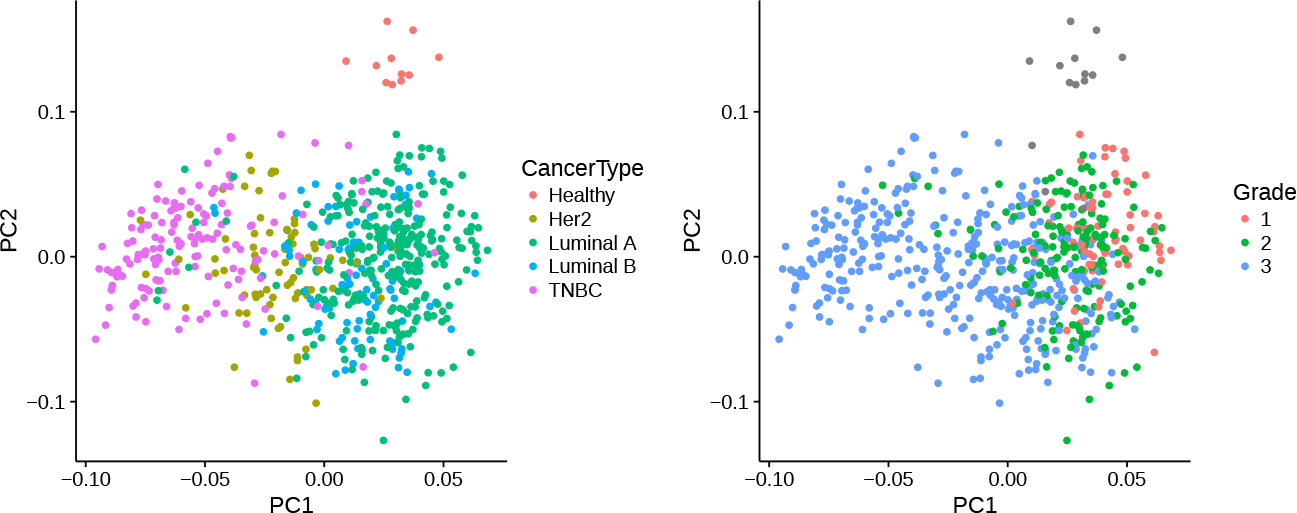
PCA plots of the breast cancer samples combined from four different microarray datasets using Maxoverall pipeline. On the left different colors designate cancer molecular subtypes, on the right — samples grade.

## Discussion

### Using RNA-seq rationale

We aimed to avoid multiple pairwise comparison of three microarray platforms described in the studies dedicated to the direct microarray data integration, which usually involves analyzing reproducibility of the genes expression rankings between different microarray datasets after applying a particular method of the probeset selection for many-to-one probesets [22]. In other words, with RNA-seq data availability we wanted to get the result unbiased towards a particular platforms design.

The similar rationale prevented us to apply the idea of using cancer samples classification based on the gene signatures with subsequent differential gene expression analysis between samples assigned to different molecular subtypes [17, 37] to determine which data processing method produces more precise classification [17, 37]. Namely, many gene signatures fail to predict outcome for a particular tumor types due to poor subtyping and in general tend to disagree with each other [55]. Moreover, Venet et al. showed that random gene signatures are significantly associated with breast cancer outcome along with the published gene signatures [56].

Therefore, in this study we used RNA-seq data as the reference for two main reasons: first, to avoid relying on the comparison of microarrays with each other, and second, to avoid using various methods of the samples classification in evaluation of microarray processing approaches, which might be a source of unreliable conclusions.

### Pipelines comparison

The reasoning behind choosing the methods of probesets selection was based on the proposed methods from the literature, which we briefly reviewed in the introduction section. Additionally, we aimed to compare probe sequence-based methods with empirical ones, based on actual probesets signals. Thus, Scores pipeline is purely based on the probe sequence characteristics and does not take into account probes signals in contrast to other methods. We chose Max pipeline as it was ranked first in Miller at al. study, while Mean pipeline is a popular approach. Finally, BrainArray approach considers both characteristics of the probes sequences and their signals as it summarizes the signals from probes mapped to a single gene.

Both Brainarray and Max pipelines performed better than other pipelines on Affymetrix and Illumina platforms, considering that Brainarray approach is not applicable to Illumina platform. Namely, Brainarray was best scored when applied to Affymetrix HuGene 1.0 ST platform, while Max performed best on the rest of platforms when comparing RNA-seq data to the individual microarray datasets. Combining datasets revealed that Brainarray pipeline correlation with RNA-seq data is slightly better than the Max pipeline. However, when merging datasets within Brainarray pipeline we had to consider Affymetrix arrays only, therefore the comparison is not strictly equivalent. Importantly, randomly picking the probeset among those mapped to same gene is the worst solution with respect to any microarray platform or pipeline used in this paper.

The task of comparing different microarray platforms to each other ideally requires using the samples from the same patients in order to ensure comparisons to be valid. Still, we consider Affymetrix HuGene 1.0 ST platform’s dynamic range being more similar to the one of RNA-seq, judging on the gene expression scatter plots for the common genes (Suppl. Fig. 10) and low-expressed genes in particular. This observation is expected as the Human Gene 1.0 ST Array utilizes a subset of the probes selected from the Exon 1.0 ST Array and focuses on a well-annotated content at the gene level, providing a more complete and more accurate picture of the gene expression than 3’-based expression array designs [28].

The obvious drawback of integrative approach is information reduction (see Table 3) as one has to choose genes properly presented in all the platforms, which might reduce the valuable information in the common genes list. However, Taminau at al. reported that merging datasets significantly increases the amount of identified differentially expressed genes in comparison to meta-analysis approach [57], which agrees with our own results, showing that direct integration of datasets from different microarray platforms improves the correlation with RNA-seq.

## Conclusions

Here we investigated five different pipelines of the microarray data processing in comparison with RNA-seq data on breast cancer samples. In particular, we aimed to assess different probesets annotations as well as different procedures of choosing probesets, which better represent the expression of their targeted genes. We show that methods, which rely on actual probesets signal intensities, are advantageous to methods considering the biological characteristics of the probes sequences only and that cross-platform integration of datasets improves correlation with the RNA-seq data. Particularly, we found that BrainArray approach based on redesigned probe-sets definition performs best on Affymetrix chips, while choosing probeset with the maximum average signal across the samples gives similar results to BrainArray on both Affymetrix and Illumina chips. Despite comparing samples from different patients, there is a strong correlation (up to 0.7) between Affymetrix arrays and RNA-seq data, while Illumina correlated worse with RNA-seq, which were not proven to be an effect of a particular transformation and normalization steps in this work. Combining datasets improved correlation with the RNA-seq data across all the pipelines and tests, proving effectiveness of the direct microarray data integration. Given that overwhelming majority of the existing and even newly uploaded data in the public databases are from microarray platforms [2], we consider the results obtained in this paper contributive to the integrative analysis as a worthwhile alternative to the classical meta-analysis of the multiple gene expression datasets.

## Author’s contributions

AF, VB, MO designed the study. AF and VB performed the data analysis. All authors read and approved the final version of the manuscript.

## Acknowledgements

We would like to express out gratitude to Prof. Louise C. Showe and her team from The Wistar Institute, Philadelphia for providing Illumina bead-level data for the GSE37614 experiment.

## Additional Files

### Additional file 1 — Supplementary Figures

This file contains quality control plots, complete set of scatter plots and correlation barplots, example of diagnostic plots for linear regression and platforms specificity and coverage scores barplots.

